# *robustica*: customizable robust independent component analysis

**DOI:** 10.1101/2021.12.10.471891

**Authors:** Miquel Anglada-Girotto, Samuel Miravet-Verde, Luis Serrano, Sarah A. Head

## Abstract

**Background:** Independent Component Analysis (ICA) allows the dissection of omic datasets into modules that help to interpret global molecular signatures. The inherent randomness of this algorithm can be overcome by clustering many iterations of ICA together to obtain robust components. Existing algorithms for robust ICA are dependent on the choice of clustering method and on computing a potentially biased and large Pearson distance matrix.

**Results:** We present *robustica*, a Python-based package to compute robust independent components with a fully customizable clustering algorithm and distance metric. Here, we exploited its customizability to revisit and optimize robust ICA systematically. From the 6 popular clustering algorithms considered, *DBSCAN* performed the best at clustering independent components across ICA iterations. After confirming the bias introduced with Pearson distances, we created a subroutine that infers and corrects the components’ signs across ICA iterations to enable using Euclidean distance. Our subroutine effectively corrected the bias while simultaneously increasing the precision, robustness, and memory efficiency of the algorithm. Finally, we show the applicability of *robustica* by dissecting over 500 tumor samples from low-grade glioma (LGG) patients, where we define a new gene expression module with the key modulators of tumor aggressiveness downregulated upon *IDH1* mutation.

**Conclusion:** *robustica* brings precise, efficient, and customizable robust ICA into the Python toolbox. Through its customizability, we explored how different clustering algorithms and distance metrics can further optimize robust ICA. Then, we showcased how *robustica* can be used to discover gene modules associated with combinations of features of biological interest. Taken together, given the broad applicability of ICA for omic data analysis, we envision *robustica* will facilitate the seamless computation and integration of robust independent components in large pipelines.

## BACKGROUND

Independent Component Analysis (ICA) is a matrix factorization method that dissects a mixture of signals into a predefined number of additive independent sources or components. ICA finds sets of statistically independent components by minimizing their mutual information^1^. In biology, ICA has a wide range of applications such as defining functional modules, removing technical noise, feature engineering, unsupervised cell type deconvolution, single-cell trajectory inference, or multi-omic analysis (reviewed in ^2^). Thanks to its information-theoretic objective function, ICA results in components that provide a simpler, more reproducible, and more biologically relevant interpretation than other popular matrix factorization methods such as Principal Component Analysis (PCA) or Non-negative Matrix Factorization (NMF)^2–7^.

*FastICA*^8^, one of the most widespread algorithms used to perform ICA, starts with a random initialization to decompose the data matrix into a source matrix and a mixing matrix of non-Gaussian independent components (Sup. Fig. 1A). In 2003, Hymberg and Hyvärinen^9^ developed *Icasso* to address the inherent randomness of *FastICA*, by running *FastICA* multiple times and clustering the components of source matrices across all runs (Sup. Fig. 1B). This clustering step involves two key choices affecting its computational efficiency and the final robust components: the distance metric and the clustering algorithm. Current implementations require pre-computing a potentially large Pearson distance square matrix to cluster components across ICA runs regardless of their different sign and order resulting from *FastICA*’s randomness^9–12^. However, correlation-based metrics are sensitive to outliers and non-Gaussian distributions as independent components, which may lead to calculating imprecise weights^13^. Additionally, more recently developed clustering algorithms could potentially improve the efficiency and quality of robust ICA.

Here, we developed *robustica*, the first Python package to carry out robust ICA with a fully customizable clustering metric and algorithm based on the powerful library *scikit-learn*^14^. By leveraging the customizability of our package to revisit and optimize the clustering step of the *Icasso* algorithm, we improved its precision, robustness, and memory efficiency. Finally, as a case study, we dissected gene expression signatures from patients with low-grade glioma (LGG) and found a set of genes related to the mechanism by which mutations in *IDH1* lead to less aggressive tumors.

## RESULTS

### *robustica* enables systematic evaluation of clustering algorithms to perform robust ICA

With *robustica*, one can fully customize the clustering method to use as long as they follow *scikit-learn* conventions^14^. We purposefully included this feature to compare how 6 different popular clustering algorithms perform at finding robust components (see Methods). As a benchmark, we dissected the >250 *E. coli* gene expression signatures from Sastry (2019)^15^ into 100 components with 100 ICA runs and selected different clustering algorithms to compute the robust components. We then evaluated the performance of the different algorithms by measuring their run time, memory usage, and silhouette scores to quantify how similar each component is to the components in the cluster compared to the components assigned to other clusters. Overall, the *DBSCAN* algorithm showed the best performance taking ∼20 seconds and ∼5500 MiB of maximum memory usage to obtain clusters with the highest median silhouette scores (*median=0*.*89*). *CommonNNClustering* also performed well, with better memory efficiency at the cost of a lower silhouette score (*median=0*.*81*) (Fig. 1B; Sup. Fig. 2-3; Sup. Tab. 1).

**Figure 1.**
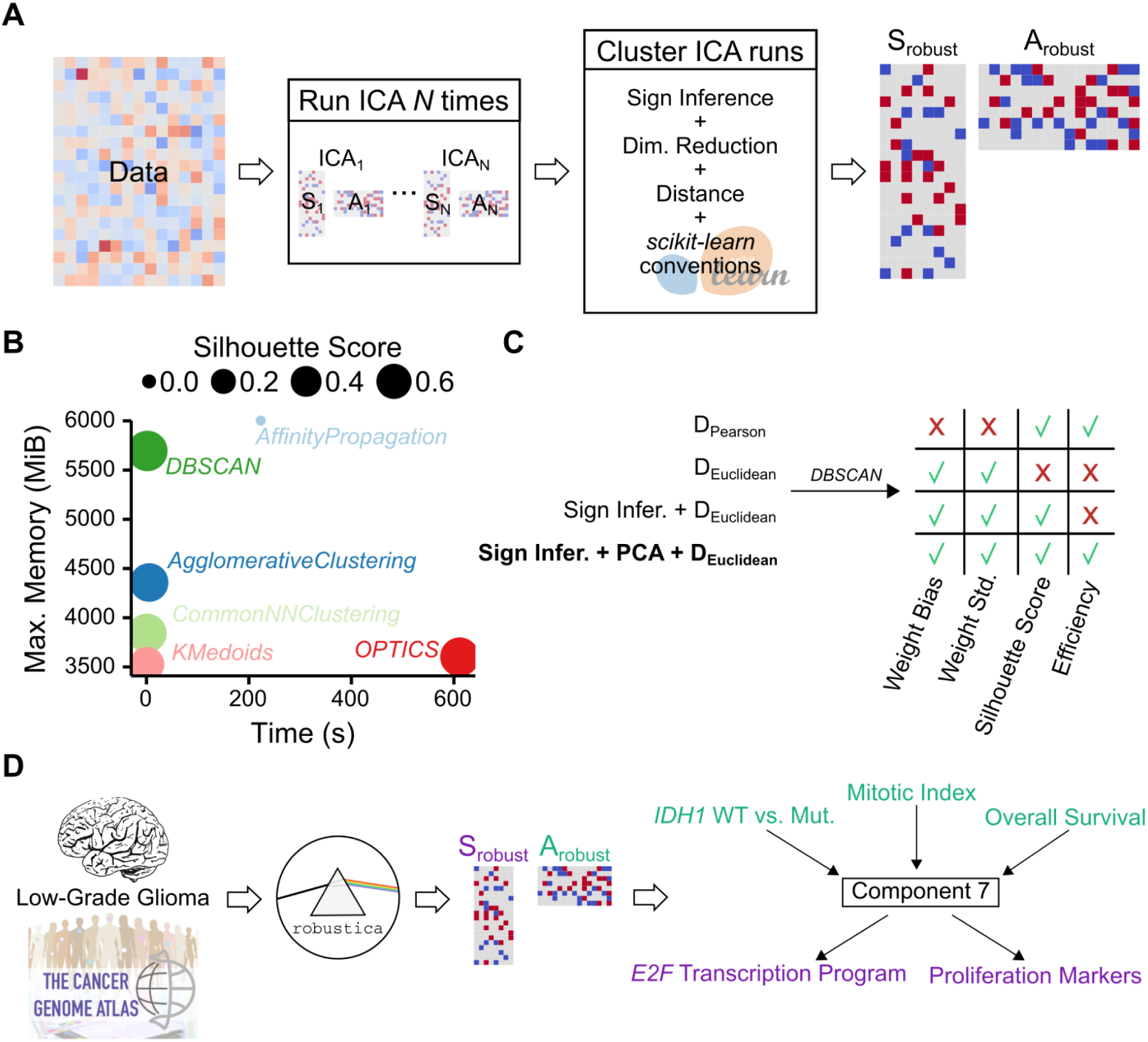
Development and implementation of *robustica* to carry out robust Independent Component Analysis (ICA). (**A**) *robustica* enables fully customized robust ICA. We built *robustica* following *scikit-learn*’s programming conventions to enable full control of both the iterative and clustering steps facilitating customization and optimization. In particular, *robustica* includes a subroutine to infer and correct the signs of components across ICA runs that improves the precision and efficiency of the clustering step by enabling us to use Euclidean distance metrics and to compress the feature space. (**B**) Comparison of clustering algorithms for robust ICA using Sastry (2019)^15^ ‘s dataset. Maximum memory usage and time for each clustering algorithm to cluster 100 ICA runs with 100 components each. Dot sizes indicate median silhouette scores (the larger the better). (**C**) Development steps to improve the precision (*Weight Std*.) and efficiency while reducing the bias of robust ICA through our sign inference-and-correction subroutine combined with PCA and Euclidean distances, using Sastry (2019)^15^ ‘s dataset. (**D**) Case study workflow for robust ICA. We dissected >500 tumor samples from LGG patients with *robustica* into 100 robust independent components. Component 7 was simultaneously associated with multiple sample features (*IDH1* mutation status, mitotic index, and overall survival) and contained genes known to be mechanistically associated with mutations in *IDH1* that modulate tumor aggressiveness.

**Supplementary Figure 1.**
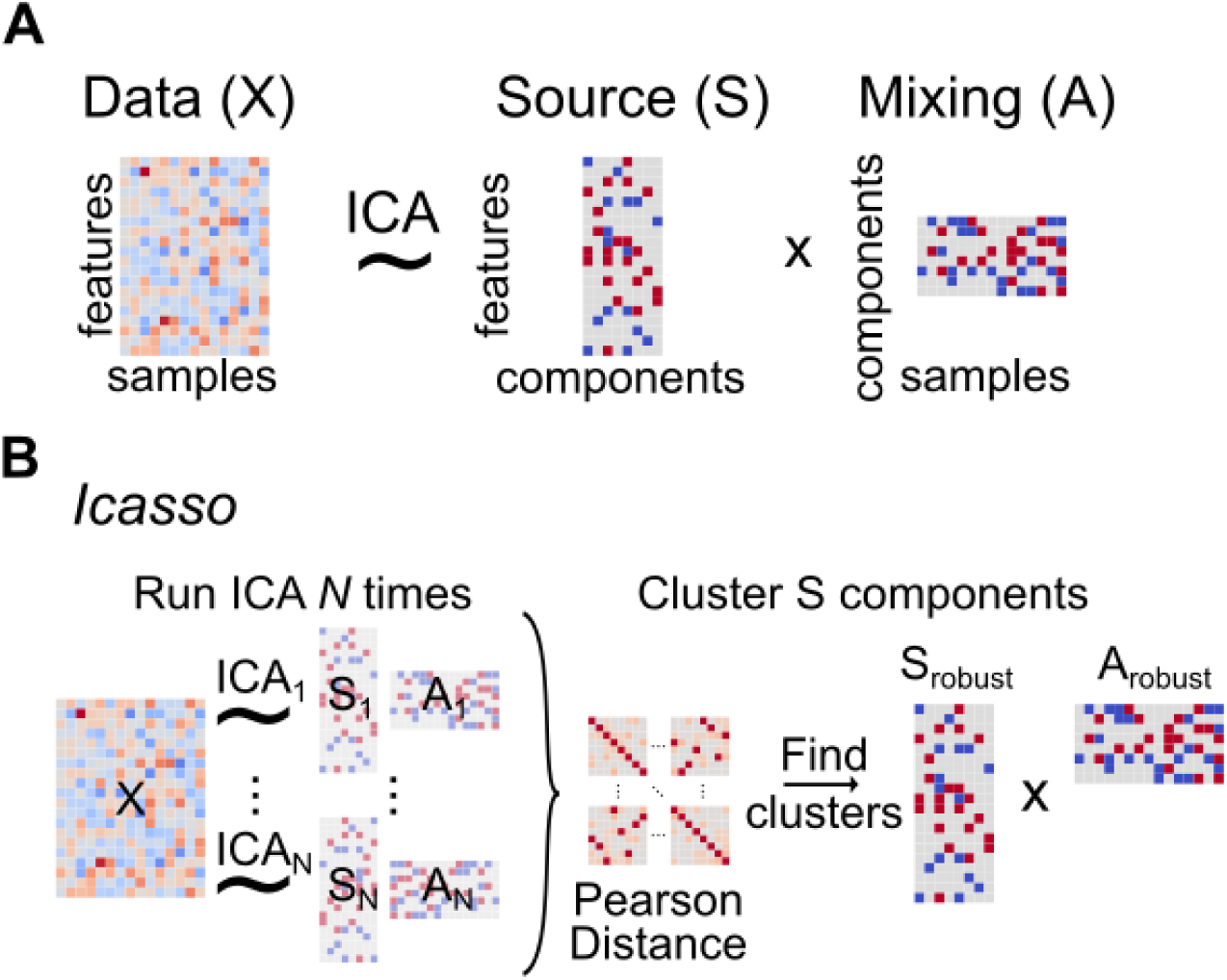
Independent Component Analysis (ICA) and robust ICA with the *Icasso* algorithm. (**A**) ICA is a matrix factorization algorithm applied for blind-source separation problems that decompose a data matrix (*X*) into *n* independent components generating a source matrix (*S*) and a mixing matrix (*A*) with information on how features and samples contribute to each independent component, respectively. (**B**) The *Icasso* algorithm overcomes the inherent randomness of the *FastICA* algorithm -a widespread algorithm to perform ICA-by running ICA multiple times and clustering the resulting independent components in *S* across all runs using an agglomerative clustering approach with average linkage and a Pearson distance matrix (*n components* · *n. runs*)^2^ dimensions as input.

**Supplementary Figure 2.**
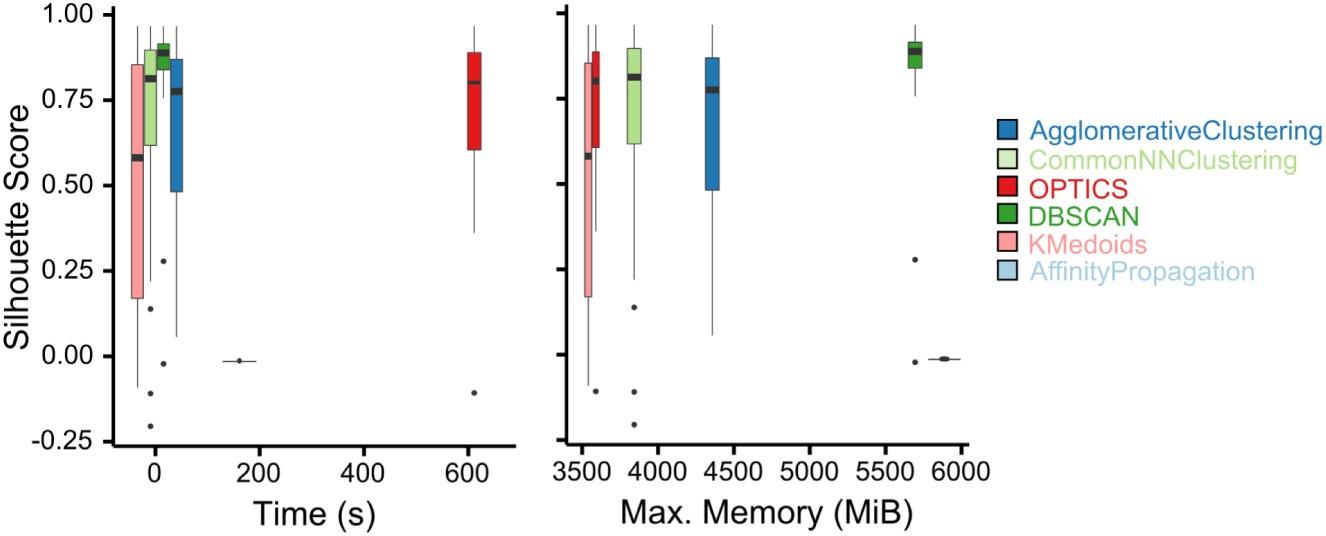
*DBSCAN* shows the best performance to compute robust independent components. Time (seconds) and maximum memory usage (MiB) compared to the average silhouette scores of each robust component (i.e. cluster) obtained using 6 different algorithms to cluster the independent components produced across 100 runs of ICA with *n_components=100* by dissecting Sastry (2019)^1^ ‘s dataset.

**Supplementary Figure 3.**
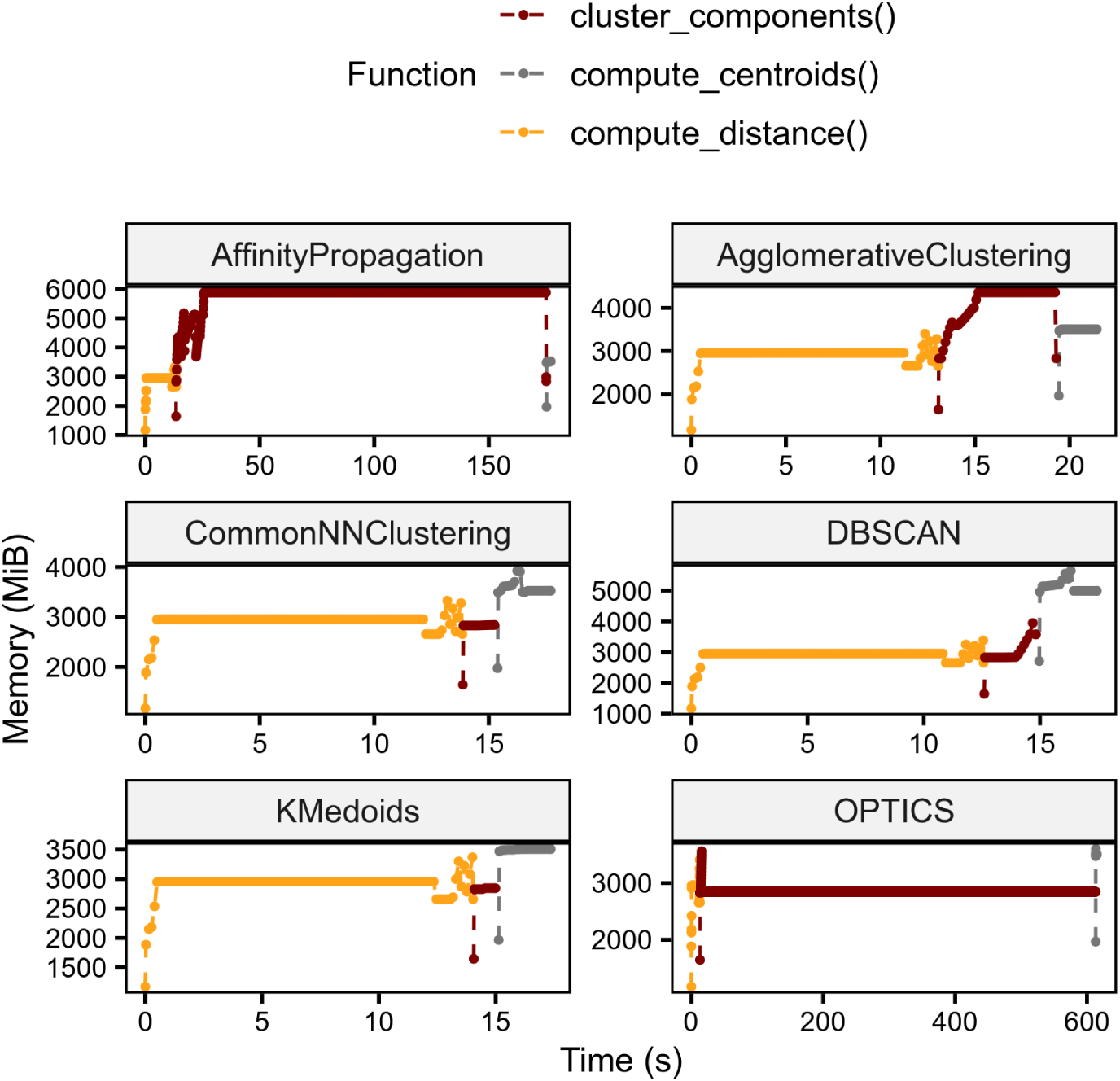
Computing the Pearson distance matrix takes the most time when using the DBSCAN algorithm. Memory usage across time and substeps (functions) to compute robust independent components in our comparison of clustering algorithms by dissecting Sastry (2019)^1^ ‘s dataset.

### Clustering ICA runs with Euclidean distances improves the precision of robust ICA

After running ICA multiple times, the *Icasso* algorithm computes a potentially large Pearson distance matrix (Sup. Fig. 1B). However, the sensitivity of Pearson distance to non-Gaussian distributions strongly biases weights in robust independent components, as the standard deviation of the weights across components in the same cluster strongly correlates with their average (Sup. Fig. 4-5). We tackled this problem with a simple subroutine to infer and correct the sign of the components across ICA runs to enable using Euclidean distances (see Methods). Our approach produced robust components with high silhouette scores and lowered the bias and standard deviation of their weights (Sup. Fig. 4-5). Since this implementation required less memory but more time to cluster sign-corrected components (Sup. Fig. 4D), we compressed the feature space of all ICA runs through PCA to reduce the overall run time and memory usage while maintaining the same performance (Sup. Fig. 4; Sup. Tab. 2; Sup. Tab. 3; Sup. Tab. 4; Sup. Tab. 5; Sup. Tab. 6).

Finally, we assessed how much the resulting gene modules differ using either Pearson or Euclidean distances (Sup. Tab. 7). While both approaches found highly similar modules, Euclidean distance tended to find more components of high silhouette scores (Sup. Fig. 6A and B). In addition, we measured how much the high precision of Euclidean distance made the modules significantly more robust to random noise compared to Pearson distance and allowed using fewer runs to recover most of the gene modules defined using 100 ICA runs to compute robust components (Sup. Fig. 6C and D; Sup. Tab. 2; Sup. Tab. 3; Sup. Tab. 4; Sup. Tab. 5; Sup. Tab. 6; Sup. Tab. 7). Bypassing the bias introduced by Pearson distances through Euclidean distances increased the reproducibility of the gene modules defined through robust ICA.

Our sign inference-and-correction subroutine creates high-quality, robust independent components by enabling us to efficiently cluster independent components across ICA runs using Euclidean distances (Fig. 1C).

**Supplementary Figure 4.**
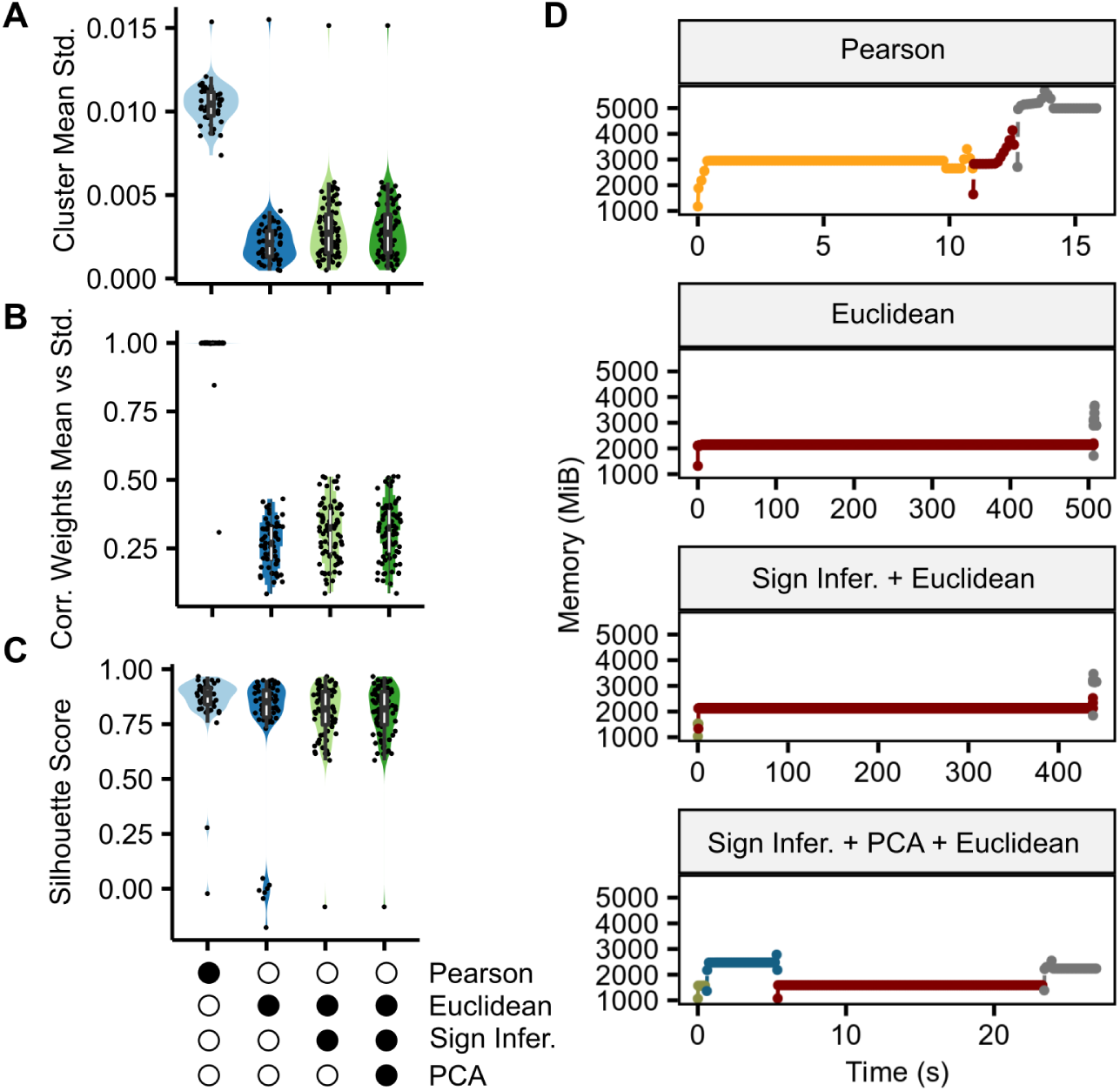
Our sign inference-and-correction subroutine combined with PCA and *DBSCAN* with Euclidean distances leads to efficiently computing unbiased and precise robust independent components. Each panel illustrates how our versions of the *Icasso* algorithm perform at dissecting Sastry (2019)^1^ ’s dataset using: the original Pearson distance matrix (Pearson); plain Euclidean distance (Euclidean); Euclidean distance after our sign inference-and-correction subroutine (Sign Infer. + Euclidean); or Euclidean distance after our sign inference-and-correction subroutine and feature compression with PCA (Sign Infer. + PCA + Euclidean). (**A**) Average standard deviations of weights used to calculate the weights of each robust independent component. (**B**) Distributions of correlations between weight averages and weight standard deviations that correspond to each robust independent component. (**C**) Average silhouette scores of each component used to calculate the robust independent components. (**D**) Memory usage across time and substeps of every version of the *Icasso* algorithm: computing Pearson distance (yellow); clustering components with *DBSCAN* (dark red); computing centroids (grey); inferring components’ signs (dark green); compressing feature space with PCA (dark blue).

**Supplementary Figure 5.**
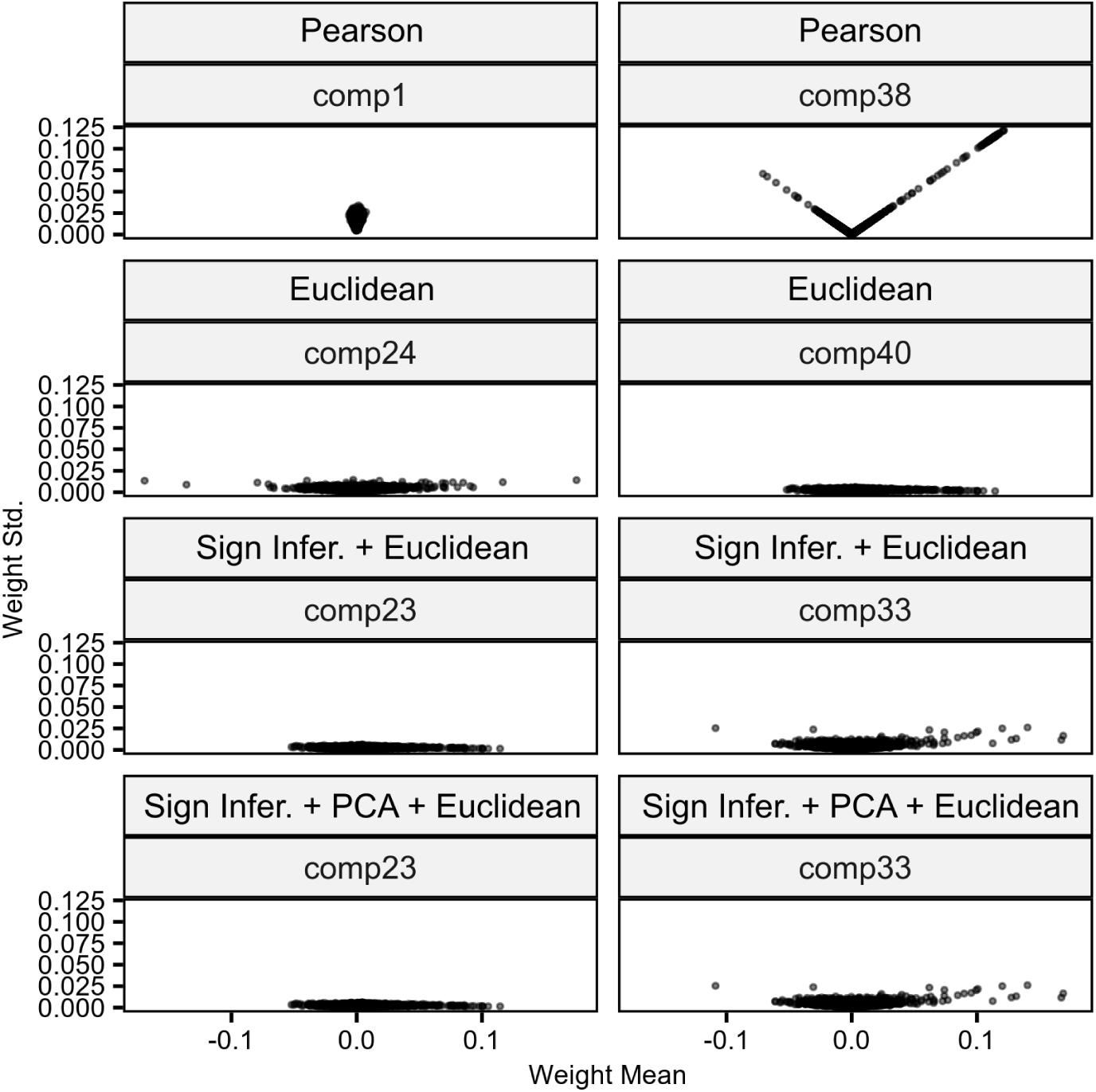
Examples of weight mean and standard deviation bias in robust independent components. The different panels illustrate how the mean and standard deviation of the components’ weights used to compute robust independent components correlated depending on which version of the *Icasso* algorithm we used to dissect Sastry (2019)^1^ ’s dataset: the original Pearson distance matrix (Pearson); plain Euclidean distance (Euclidean); or Euclidean distance after our sign inference-and-correction subroutine (Sign Infer. + Euclidean); Euclidean distance after our sign inference-and-correction subroutine and feature compression with PCA (Sign Infer. + PCA + Euclidean).

**Supplementary Figure 6.**
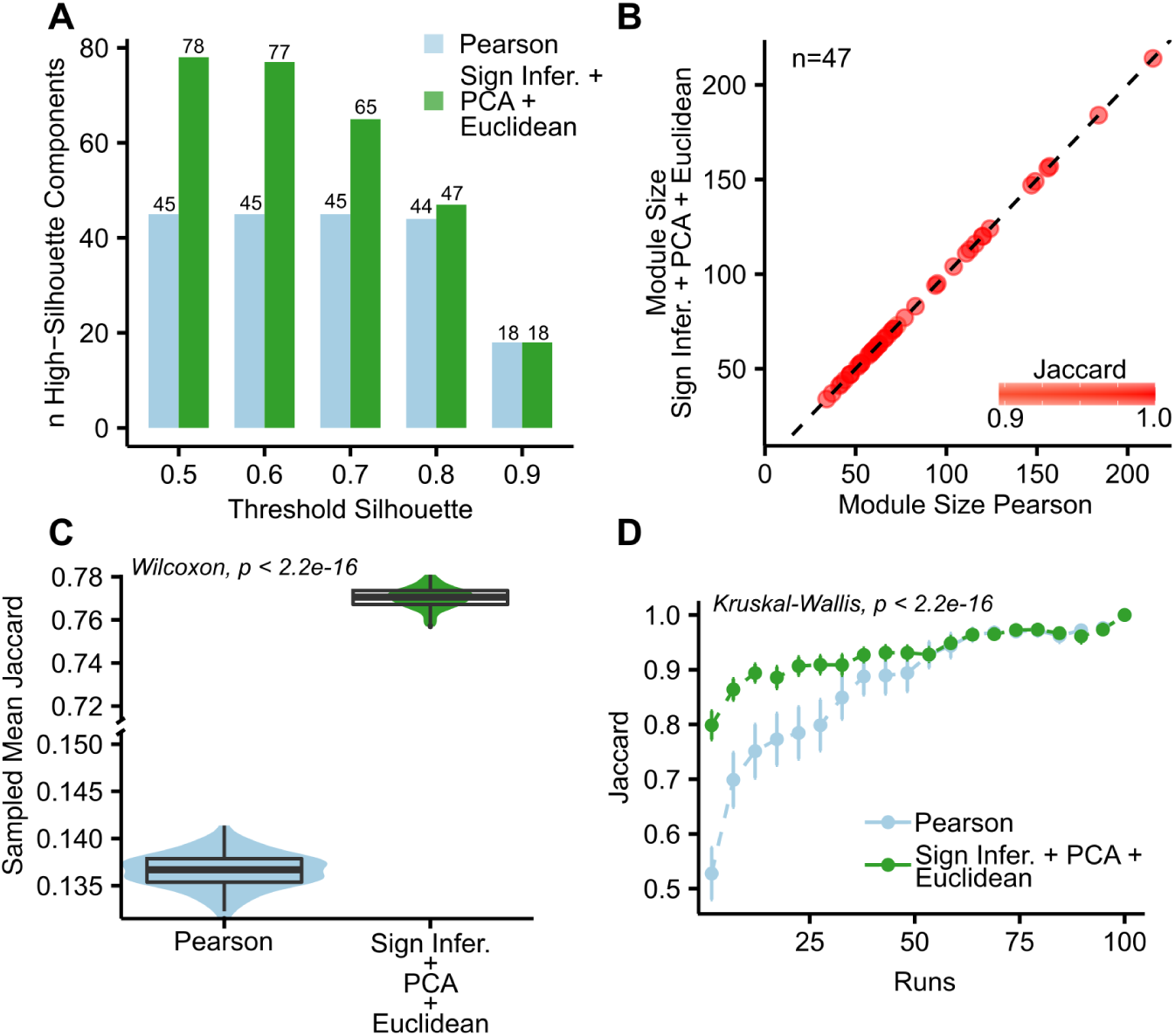
Comparison between gene modules obtained from Sastry (2019)^1^ ’s dataset with the classical (Pearson) and the revisited (Sign Infer. + PCA + Euclidean) *Icasso* algorithm to perform robust ICA. (**A**) Number of components with high silhouette scores using different thresholds. (**B**) Sizes of gene modules uniquely mapped between the two approaches (47 in total). The color gradient indicates the degree of Jaccard similarity between mapped modules. (**C**) Average Jaccard similarities between the gene modules defined using inferred robust independent components and their corresponding weights subjected to random noise sampled 100 times. (**D**) Distributions (mean and standard deviation) of maximum Jaccard similarities between the gene modules defined using a different number of ICA runs compared with the gene modules defined using 100 ICA runs for both the classical and the revisited approaches.

### *robustica* recovers a gene expression module with the key regulators of tumor aggressiveness in LGG mechanistically associated to mutations in *IDH1*

As a case study, we dissected gene expression profiles from >500 LGG tumor samples from The Cancer Genome Atlas (TCGA). LGGs are characterized by mutations in the isocitrate dehydrogenase (*IDH1*) enzyme that decrease tumor aggressiveness by indirectly inhibiting the *E2F* transcription program, an important switch controlling homeostasis and tumorigenesis^16–18^. We then explored whether certain gene modules associated with *IDH1* mutations recovered known molecular mechanisms.

Weights in component 7 were highly associated with *IDH1* mutation status, survival probability, and expression-based indices of cell proliferation (Sup. Fig. 7A-C). We related these sample traits to the gene expression signatures by defining a module of 421 genes (Sup. Fig. 7D; Sup. Tab. 8) which, as expected, was enriched in proliferation-related biological processes (Sup Fig. 8; Sup. Tab. 8) and contained 7 out of the 9 genes used to compute the mitotic index^19^ and known proliferation markers as *MKI67*^20^. *Interestingly, our gene module also included 4 E2F* transcription factors (*E2F1, E2F2, E2F7, E2F8*) and was enriched with 98 targets of *E2F*s (Sup. Fig. 9; Sup. Tab. 8).

With this, we demonstrate the utility of *robustica* to identify gene sets whose combined expression in LGG is associated with genotypes and phenotypes of interest. Our identified module contained downstream effectors controlling cell proliferation associated with *IDH1* mutation status and tumor aggressiveness (Fig. 1D), demonstrating the biological applicability of this approach.

**Supplementary Figure 7.**
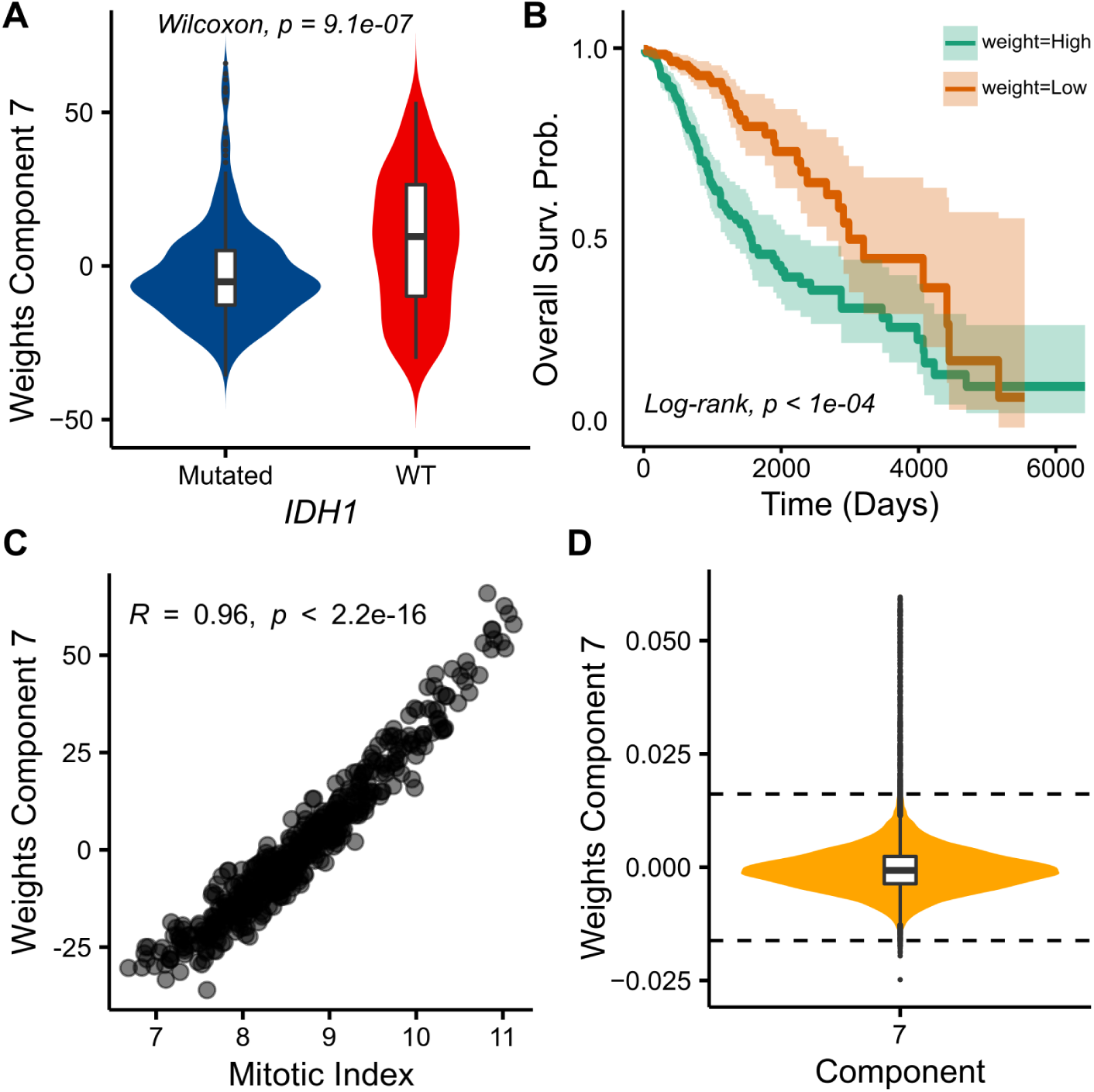
Independent component 7 defines a module simultaneously associated with LGG patients’ mutation status in *IDH1*, survival probability, and mitotic index. (**A**) Distribution of sample weights in component 7 in the robust mixing matrix among patients with (Mutated) or without (WT) mutations in gene *IDH1*. We used a Wilcoxon Rank Sum test to assess the statistical differences between the groups (top label). (**B**) Kaplan-Meier curves of patients with weights higher or lower than the median weight in component 7 in the robust mixing matrix. The p-value of the log-rank test between the two groups is indicated at the bottom left. (**C**) Relationship between weights in component 7 in the robust mixing matrix and each sample’s mitotic index. The Spearman correlation coefficient and its corresponding p-value are indicated at the top left (*R* and *p*, respectively). (**D**) Distribution of weights in component 7 in the robust source matrix used to define the gene module corresponding to our sample features of interest. The horizontal dashed lines indicate the thresholds applied to define the module.

**Supplementary Figure 8.**
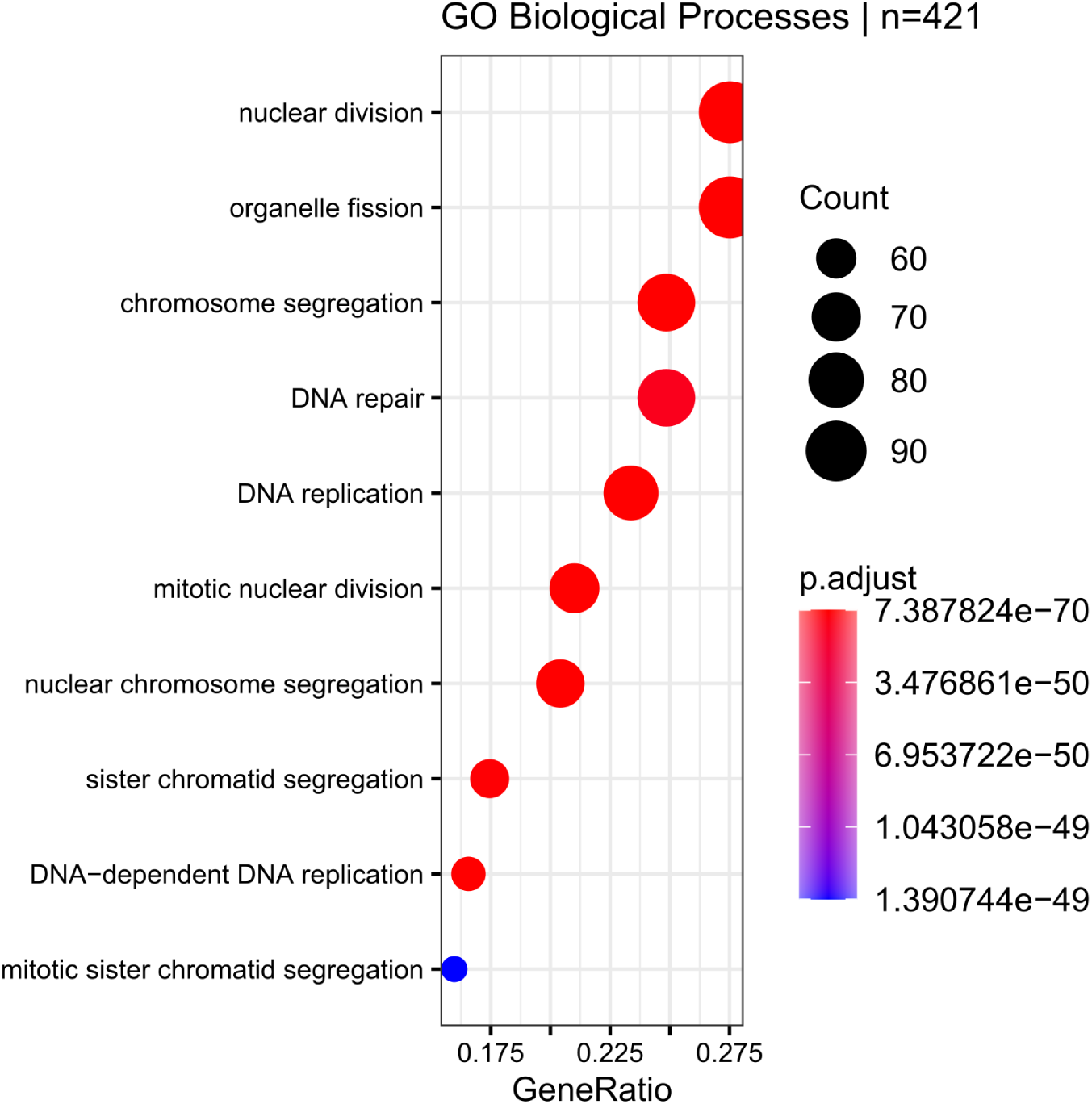
The gene module defined from component 7 is enriched in proliferation-related biological processes. Results from overrepresentation tests of our gene module defined from component 7 and Gene Ontology (GO) Biological Processes.

**Supplementary Figure 9.**
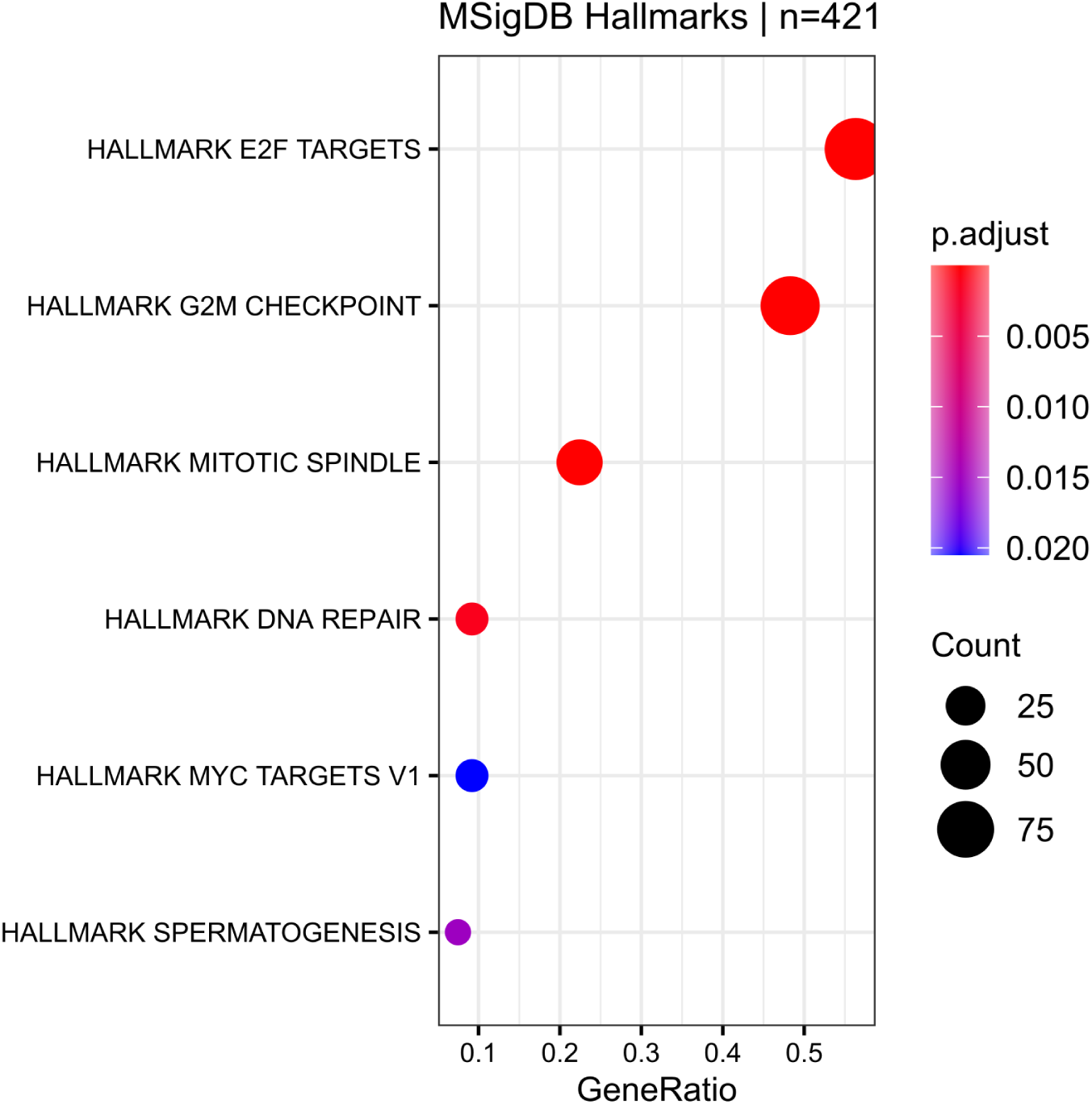
The gene module defined from component 7 is enriched in targets of *E2F* transcription factors, known modulators of glioma aggressiveness. Results from overrepresentation tests of our gene module defined from component 7 and MSigDB Hallmarks.

## CONCLUSION

We created *robustica*, a new Python package built on top of *scikit-learn* that enables performing precise, efficient, and customizable robust ICA seamlessly. Using different clustering algorithms and distance metrics, we tested whether robust ICA could be further optimized. Our sign correction subroutine improved the precision, robustness, and memory efficiency of the clustering step. As an example, we explored how the gene modules generated with *robustica* from transcriptomic profiles of LGG patients are associated with multiple markers of the disease’s aggressiveness simultaneously. Overall, *robustica* makes high-performance robust ICA more accessible to potentially analyze any omic data type and facilitates its incorporation in larger computational pipelines.

## IMPLEMENTATION

*robustica* is written in Python under the open-source BSD 3-Clause license. The source code and documentation are freely available at https://github.com/CRG-CNAG/robustica. Additionally, all scripts to reproduce the work presented are available at https://github.com/MiqG/publication_robustica.

The algorithm to carry out robust ICA is controlled by the main class *RobustICA*. After instantiation, one can use the *fit* method on a data matrix to run the *FastICA* algorithm multiple times and cluster the resulting independent components with the desired number of components and clustering distance metric and algorithm. Subsequently, one can recover the computed source and mixing matrices through the *transform* method.

To facilitate the customizability and seamless integration of the algorithm, the user can specify all the parameters related to every step of the robust ICA algorithm as similarly as possible to the core *FastICA* class already available in *scikit-learn*. For this reason, we only required 6 more arguments labeled using the “robust_” prefix. These arguments determine: the number of times to run *FastICA* before clustering (*robust_runs*), whether to use our subroutine to infer and correct the signs of the components across *FastICA* runs (*robust_infer_signs*), the custom clustering algorithm class (*robust_method*), the keywords to pass to the clustering algorithm class (*robust_kws*), whether to speed up the clustering step using PCA to reduce the dimensions of the independent components across all runs (*robust_dimreduce*) and, when the distance needs to be precomputed, the function to use to compute pairwise distances in the clustering step (*robust_precompdist_func*).

## AVAILABILITY OF DATA AND MATERIALS

- source code and documentation: https://github.com/CRG-CNAG/robustica
- analysis pipeline in this paper: https://github.com/MiqG/publication_robustica

## AVAILABILITY AND REQUIREMENTS

**Project name:** *robustica*

**Project home page:** https://github.com/CRG-CNAG/robustica

**Operating system(s):** Platform Independent.

**Programming language:** Python.

**Other requirements:** *numpy, pandas, scikit-learn, scikit-learn-extra, joblib, tqdm*.

**License:** BSD 3-Clause.

**Any restrictions to use by non-academics:** None.

## LIST OF ABBREVIATIONS

ICA: Independent Component Analysis
LGG: Low-Grade Glioma
PCA: Principal Component Analysis
NMF: Non-negative Matrix Factorization
TCGA: The Cancer Genome Atlas
IDH1: isocitrate dehydrogenase

## METHODS

All scripts to reproduce the work presented are available at https://github.com/MiqG/publication_robustica.

### Data download and preprocessing

We downloaded the 278 gene expression signatures for 3’923 genes in *E. coli* from Sastry (2019)^1^ ‘s Supplementary Data 1. Following the author’s procedure, we normalized the expression signatures with respect to controls (“control wt_glc 1” and “control wt_glc 2”) by subtracting the mean gene expression of the controls to the rest of the signatures.

We used the XenaBrowser^2^ to obtain the pan-cancer log-normalized gene expression signatures (530 samples and 20’531 genes), somatic mutation, and clinical information in tumor samples from low-grade glioma (LGG) patients in The Cancer Genome Atlas (TCGA). We defined *IDH1* mutated samples as those with at least one missense or nonsense mutation in the gene.

We computed the mitotic index for each sample by averaging the log-normalized expression levels of 9 genes (*CDKN3, ILF2, KDELR2, RFC4, TOP2A, MCM3, KPNA2, CKS2*, and *CDK1*) as described in Yang (2016)^3^.

### Comparing different clustering algorithms to compute robust independent components

We used the preprocessed *E. coli* dataset to compare how different clustering algorithms perform in terms of computing time, memory usage and cluster average silhouette scores, and cluster weight standard deviation. We followed the *Icasso* procedure described in Himberg and Hyvärinen (2003)^4^ and modified the clustering step. First, we standardized the gene expression matrix across genes (rows). Then, we used *robustica* to run *FastICA* 100 times with 100 components and default parameters. We saved the source (S) and mixing (A) matrices generated across the runs to ensure that we used the same inputs for the clustering step. Subsequently, we created a new instance of *robustica* to cluster the components across the S matrices using the precomputed distance matrix based on absolute Pearson correlations as input:

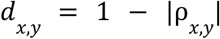

where *ρ*_*x, y*_ is the Pearson correlation between components *x* and *y* and *d*_*x, y*_ is the distance between components *x* and *y* which generates a square matrix of (*n. components* · *n. runs*) rows and (*n. components* · *n. runs)* columns.

We used the following clustering algorithms from the *scikit-learn* and *scikit-learn-extra* libraries that accepted a precomputed dissimilarity matrix as input:

- *sklearn*.*cluster*.*AgglomerativeClustering* with *linkage=”average”* and *n_clusters=100*.
- *sklearn*.*cluster*.*AffinityPropagation* with *n_clusters=100*.
- *sklearn*.*cluster*.*DBSCAN* with *min_samples=50*.
- *sklearn*.*cluster*.*OPTICS* with *min_samples=50*.
- *sklearn_extra*.*cluster*.*KMedoids* with *n_clusters=100*.
- *sklearn_extra*.*cluster*.*CommonNNClustering* with *min_samples=50*.

Finally, we profiled the run time and memory usage of the algorithms and we evaluated their performance based on the distribution of average silhouette scores per cluster through *sklearn*.*metrics*.*silhouette_samples* with the default parameters (Sup. Tab. 1).

### Inference of components’ signs across multiple ICA runs

To be able to cluster ICA components across different runs regardless of their sign and order without precomputing the correlation-based dissimilarity matrix, we equipped *robustica* with a subroutine that infers and corrects the signs of the components before the clustering step. After running *FastICA* multiple times, the subroutine sets the first run of ICA as a reference and computes the pairwise Pearson correlation between the reference and every other run. For every pair, we obtain a correlation matrix and, assuming that every component will only have one similar component in another run since ICA returns independent components, we store the sign of the maximum absolute correlation indicating whether the sign has changed with respect to the reference and use it to correct the signs of the components across runs.

### Module definition

Throughout this article, we defined gene modules as described in Saelens (2018)^5^. For every component in the source matrix, we used *fdrtool::fdrtool* to estimate each gene’s false discovery rate (FDR) based on the distribution of weights and applied a cutoff of FDR<0.01 to select the genes belonging to every module.

### Subjecting robust components to random noise to quantify their robustness

Robust independent components inferred using Pearson distance showed a bias between the mean weight magnitude and weight standard deviation across ICA runs (Sup. Fig. 4 and 5). We measured how this uncertainty affects the resulting gene modules when using either Pearson or Euclidean distances with sign-corrected components. We used *np*.*random*.*normal* to draw 100 samples of weights of robust independent components using their own mean and standard deviation. Then, for each sample, we defined the gene modules as described above and measured how much the sampled modules differed from the original modules using Jaccard similarity with 1 - *sklearn*.*metrics*.*pairwise_distances(metric=”jaccard”)*. We used the maximum Jaccard similarity to map the pairs of modules to each other. Finally, we saved summary statistics of the mapped pairs (Sup. Fig. 6C, Sup. Tab. 2).

### Mapping modules between *robustica* outputs

We compared how the gene modules resulting from our sign-corrected feature-compressed robust ICA with Euclidean distance metrics differed from the gene modules resulting from using the classical *Icasso* algorithm (Sup. Tab. 7) by measuring pairwise Jaccard similarities using *proxy::sim*, resulting in a square matrix.

We then used module similarities to map the gene modules from one procedure to the other by taking the pair with maximum Jaccard similarity.

### Exploring LGG expression profiles through robust ICA

We used *robustica* to dissect the LGG expression profiles into robust independent components using our revisited *Icasso* algorithm (sign inference + PCA + Euclidean distance) with 100 components across 100 ICA runs (Sup. Tab. 8). We did not consider those components with an average silhouette score lower than 0.9.

Then, we explored which components in the mixing matrix were associated with *IDH1* mutation status through an unpaired Wilcoxon Rank Sum test (*stats::wilcox*.*test*); with patient overall survival through a Cox Proportional-Hazards regression (*survival::coxph*); and with the mitotic index through Spearman correlation (*stats::cor*).

Finally, we further characterized the gene signatures in the source matrix that drive these associations in component 7. We defined a gene set by taking the genes with extreme weights in that component as explained above and performed a gene set enrichment analysis for biological processes (*clusterProfiler::enrichGO*) and MSigDB’s hallmark signatures (*clusterProfiler::enricher*) using the default parameters.

### Software

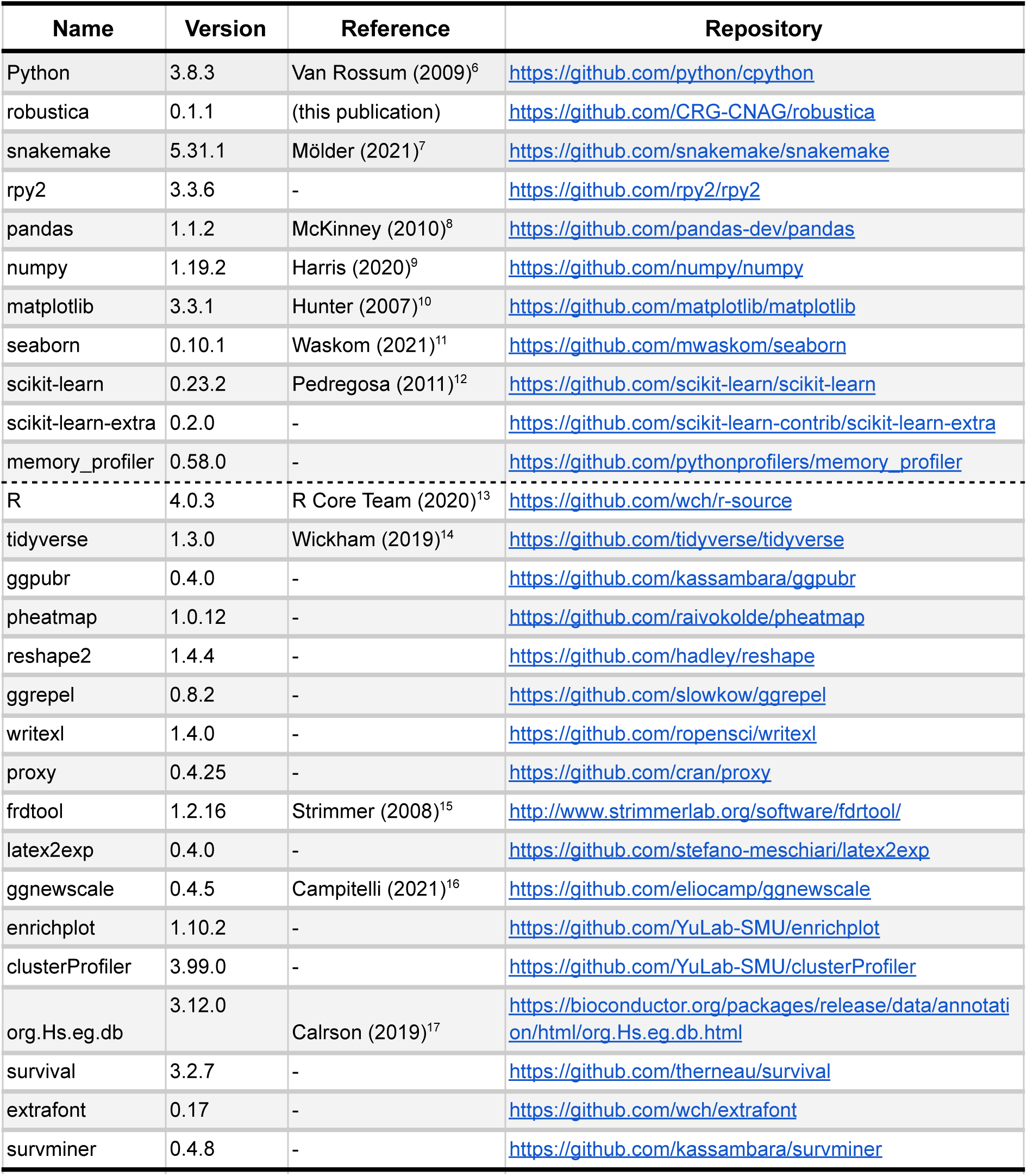

## SUPPLEMENTARY TABLES

**Supplementary Table 1. Evaluation of 6 different algorithms in clustering 100 independent components from 100 runs of ICA into robust independent components**. Memory usage profiles across time for each subroutine of the clustering step for each of the algorithms considered (*performance_evaluation*); the clustering evaluation for each of the components for each of the algorithms considered (*clustering_evaluation*); and overall summary stats of the clustering evaluation (*clustering_evaluation_summary*).

**Supplementary Table 2. Evaluation of different versions of the *Icasso* algorithm to perform robust ICA using Pearson distances (*icasso*), Euclidean distances (*robustica_nosign*), our subroutine for component sign inference and correction together with Euclidean distances (*robustica*), and our subroutine for component sign inference and correction and compression of the feature space with PCA together with Euclidean distances (*robustica_pca*)**.

Memory usage (MiB) profiles across time (seconds) for each version of the *Icasso* algorithm (*performance_evaluation)*. Clustering scores of the components for each version of the algorithm (*clustering_evaluation*). Evaluation of gene module reproducibility using a different number of ICA runs to compute robust independent components for each version of the algorithm (*module_mapping_diff_runs*). Evaluation of gene module robustness upon subjecting robust independent components estimated with the different versions of the *Icasso* algorithm to random noise (*module_mapping_robustness*). Pairwise Jaccard similarity between gene modules defined from using two different versions of the *Icasso* algorithm: the original Pearson distance matrix (rows) and Euclidean distance after our sign inference-and-correction subroutine and feature compression with PCA (columns) (*gene_modules_jaccard_similarity*). Pairs of modules mapped with Jaccard similarities and their corresponding features (*gene_modules_mapping*).

**Supplementary Table 3. Source matrix weight means from Sastry (2019)^1^**.

Source matrix weight means computed with different versions of the *Icasso* algorithm using Pearson distances (*icasso*), Euclidean distances (*robustica_nosign*), our subroutine for component sign inference and correction together with Euclidean distances (*robustica*), and our subroutine for component sign inference and correction and compression of the feature space with PCA together with Euclidean distances (*robustica_pca*).

**Supplementary Table 4. Source matrix weight standard deviation from Sastry (2019)^1^**. Source matrix weight standard deviation computed with different versions of the *Icasso* algorithm using Pearson distances (*icasso*), Euclidean distances (*robustica_nosign*), our subroutine for component sign inference and correction together with Euclidean distances (*robustica*), and our subroutine for component sign inference and correction and compression of the feature space with PCA together with Euclidean distances (*robustica_pca*).

**Supplementary Table 5. Mixing matrix weight means from Sastry (2019)**^1^.

Mixing matrix weight means computed with different versions of the *Icasso* algorithm using Pearson distances (*icasso*), Euclidean distances (*robustica_nosign*), our subroutine for component sign inference and correction together with Euclidean distances (*robustica*), and our subroutine for component sign inference and correction and compression of the feature space with PCA together with Euclidean distances (*robustica_pca*).

**Supplementary Table 6. Mixing matrix weight standard deviation from Sastry (2019)^1^**. Mixing matrix weight standard deviation computed with different versions of the *Icasso* algorithm using Pearson distances (*icasso*), Euclidean distances (*robustica_nosign*), our subroutine for component sign inference and correction together with Euclidean distances (*robustica*), and our subroutine for component sign inference and correction and compression of the feature space with PCA together with Euclidean distances (*robustica_pca*).

**Supplementary Table 7. Gene modules defined from Sastry (2019)**^1^.

Gene modules represented as binary matrices defined for each version of the *Icasso* algorithm using Pearson distances (*icasso*), Euclidean distances (*robustica_nosign*), our subroutine for component sign inference and correction together with Euclidean distances (*robustica*), and our subroutine for component sign inference and correction and compression of the feature space with PCA together with Euclidean distances (*robustica_pca*).

**Supplementary Table 8. Exploring LGG expression profiles through robust ICA**.

This file contains robust source (*source_matrix_S*) and mixing (*source_matrix_A*) matrices resulting from running robust ICA with 100 ICA runs and 100 components with *robustica*. And sample information: the sample metadata (*sample_metadata*), mitotic index (*sample_indices*). We also included the results from our analysis for all robust independent components (*mutation_association, survival_association, corr_sample_indices, gene_modules*), and the gene-set enrichment results for the gene module defined from component 7 (*enrichment-GO_BP, enrichment-MSigDB_Hallmarks*).

## AUTHOR CONTRIBUTIONS

MAG created the software. SMV, LS, SH supervised the project. All authors contributed to designing the analysis and writing the manuscript.

## ACKNOWLEDGEMENTS

We thank Xavier Hernandez-Alias, Marc Weber, and Leandro G. Radusky for the insightful discussions throughout the development of this project.

The results shown here are in part based upon data generated by the TCGA Research Network: https://www.cancer.gov/tcga.

## REFERENCES

1. Herault, J. & Ans, B. Réseau de neurones à synapses modifiables: décodage de messages sensoriels composites par apprentissage non supervisé et permanent. Réseau Neurones À Synap. Modif. Décodage Messag. Sensoriels Compos. Par Apprentiss. Non Supervisé Perm. 299, 525–528 (1984).

2. Sompairac, N. et al. Independent Component Analysis for Unraveling the Complexity of Cancer Omics Datasets. Int. J. Mol. Sci. 20, (2019).

3. Liebermeister, W. Linear modes of gene expression determined by independent component analysis. Bioinformatics 18, 51–60 (2002).

4. Lee, S.-I. & Batzoglou, S. Application of independent component analysis to microarrays. Genome Biol. 4, R76 (2003).

5. Stein-O’Brien, G. L. et al. Enter the Matrix: Factorization Uncovers Knowledge from Omics. Trends Genet. 34, 790–805 (2018).

6. Cantini, L. et al. Assessing reproducibility of matrix factorization methods in independent transcriptomes. Bioinformatics 35, 4307–4313 (2019).

7. Way, G. P., Zietz, M., Rubinetti, V., Himmelstein, D. S. & Greene, C. S. Sequential compression of gene expression across dimensionalities and methods reveals no single best method or dimensionality. bioRxiv 573782 (2019) doi:10.1101/573782.

8. Hyvärinen, A. & Oja, E. A Fast Fixed-Point Algorithm for Independent Component Analysis. Neural Comput. 9, 1483–1492 (1997).

9. Himberg, J. & Hyvarinen, A. Icasso: software for investigating the reliability of ICA estimates by clustering and visualization. in 2003 IEEE XIII Workshop on Neural Networks for Signal Processing (IEEE Cat. No.03TH8718) 259–268 (2003). doi:10.1109/NNSP.2003.1318025.

10. Biton, A. MineICA: Analysis of an ICA decomposition obtained on genomics data. (Bioconductor version: Release (3.13), 2021). doi:10.18129/B9.bioc.MineICA.

11. LabBandSB/BIODICA: ‘Independent Component Analysis of BIg Omics Data’. GitHub https://github.com/LabBandSB/BIODICA.

12. Zheng, C.-H., Huang, D.-S., Kong, X.-Z. & Zhao, X.-M. Gene Expression Data Classification Using Consensus Independent Component Analysis. Genomics Proteomics Bioinformatics 6, 74–82 (2008).

13. Jiang, D., Tang, C. & Zhang, A. Cluster analysis for gene expression data: a survey. IEEE Trans. Knowl. Data Eng. 16, 1370–1386 (2004).

14. Pedregosa, F. et al. Scikit-learn: Machine Learning in Python. J. Mach. Learn. Res. 12, 2825–2830 (2011).

15. Sastry, A. V. et al. The Escherichia coli transcriptome mostly consists of independently regulated modules. Nat. Commun. 10, 5536 (2019).

16. Miyata, S. et al. An R132H Mutation in Isocitrate Dehydrogenase 1 Enhances p21 Expression and Inhibits Phosphorylation of Retinoblastoma Protein in Glioma Cells. Neurol. Med. Chir. (Tokyo) 53, 645–654 (2013).

17. Youssef, G. & Miller, J. J. Lower Grade Gliomas. Curr. Neurol. Neurosci. Rep. 20, 21 (2020).

18. Fang, Z. H. & Han, Z. C. The transcription factor E2F: a crucial switch in the control of homeostasis and tumorigenesis. Histol. Histopathol. 403–413 (2006) doi:10.14670/HH-21.403.

19. Yang, Z. et al. Correlation of an epigenetic mitotic clock with cancer risk. Genome Biol. 17, 205 (2016).

20. Gerdes, J. et al. Cell cycle analysis of a cell proliferation-associated human nuclear antigen defined by the monoclonal antibody Ki-67. J. Immunol. 133, 1710–1715 (1984).

